# A Mechanical time window for correct proprioceptor development

**DOI:** 10.64898/2026.10.24.761569

**Authors:** Abeer Hassan, Naomi Halachmi, Adi Salzberg, Shelly Tzlil

**Affiliations:** Faculty of Mechanical Engineering, Technion – Israel Institute of Technology, Haifa 3200003 Israel; Department of Genetics and Developmental Biology, The Rappaport Faculty of Medicine and Research Institute, Technion-Israel Institute of Technology, Haifa 3109601, Israel

## Abstract

Proprioception requires the transduction of mechanical strain generated by muscle contractions into sensory neuron activity. Although mechanical forces are essential for the *function* of mechanosensory organs, their role in the *development* of these organs remains largely unknown. It was already shown that the fly chordotonal organ (ChO), a multicellular proprioceptor, must acquire correct mechanical properties for proper function. Here, using an *in vivo* approach, we identify a critical ‘mechanical’ time window during which the ChOs must experience appropriate mechanical forces to acquire the mechanical properties required for sensory function.

To define this developmental window, we downregulated Pericardin (Prc), a major component of the ChO extracellular matrix, at different time points during embryonic and larval development. We then measured *in vivo* the efficiency with which muscle-generated deformations are converted into compressive strain in the ChO cap cell, a key step in transmitting mechanical forces to the sensory neuron. The most severe effect on conversion efficiency was evident when *prc* was downregulated during late embryogenesis, a time window which coincides with the onset of embryonic muscle contractions. This observation suggests that muscle-generated forces provide developmental cues required for the development of ChOs with correct mechanical properties. Consistent with this hypothesis, reducing muscle-generated forces during the same embryonic period resulted in a similar phenotype of impaired ChO mechanics and sensory function. Together, our findings reveal a previously unknown role for muscle-generated forces in shaping the development of fully functional proprioceptors.

## Introduction

Proprioception is the ability to sense the posture and movement of body parts based on signals from within the body. It requires the transduction of mechanical strain generated by muscle contractions into sensory neuron impulses via the activation of mechano-sensitive ion channels. Clearly, mechanical signals are essential for the *function* of these mechano-sensitive organs, however, the role of mechanical signals in their *development* is currently largely unknown.

Invertebrate and vertebrate proprioceptors are structured differently; however, the underlying principle of their action is similar. In both systems, a sensory neuron responds to mechanical deformations generated by muscle contractions via stretch-sensitive ion-channels. Muscle-generated deformations propagate to the sensory neuron through a connective structure composed of accessory cells and an extracellular matrix (ECM).

In mammals, a prominent proprioceptive input is provided by muscle spindles, whereas in invertebrates, proprioception is largely mediated through specific subclasses of multiple dendritic neurons ^1–3^ and stretch-responsive multicellular proprioceptors that are named chordotonal organs (ChO) because of their chord-like shape ^4^.

The ChO is structured as a linear array of six different cell types. The sensory neuron, which is enwrapped by a glia cell (scolopale), is stretched between two accessory cells called *cap* and *ligament*, which in turn are attached to the cuticle (chitin-rich ECM constituting the exoskeleton) via specialized attachment cells (Figure 1a). Muscles pull on the cuticle, whose deformation is transmitted to the sensory neuron via the accessory cells and the associated ECM (i.e., the neuron senses muscle contractions indirectly). Thus, cap and ligament cells need to develop appropriate mechanical properties to transmit the correct information to the neuron.

**FIGURE 1:**
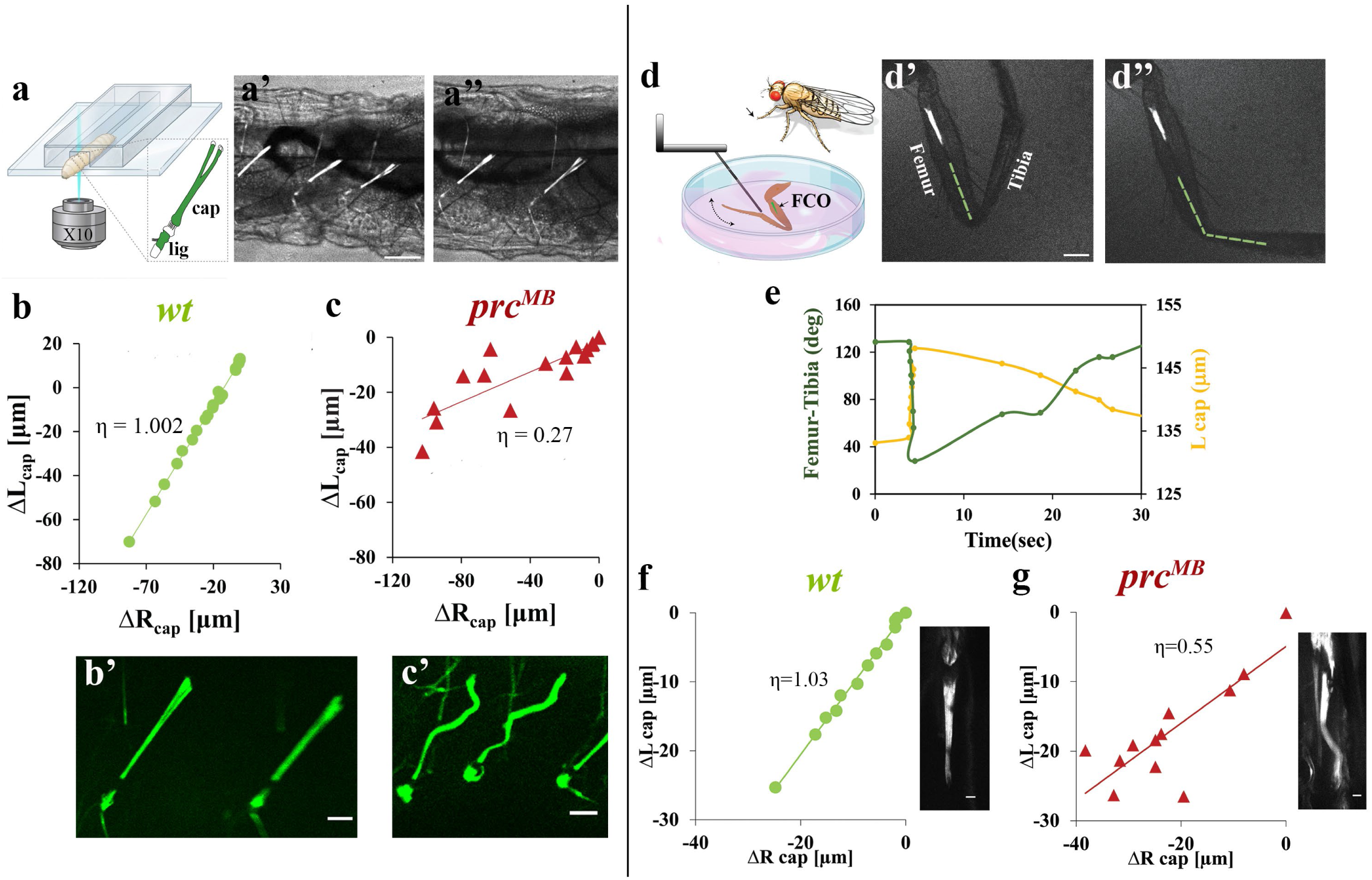
Sensory organ function is compromised in larvae and adult flies with modified mechanical properties: **(a-a”)**: Illustration of the experimental setup for larva crawling in an agarose channel. Agarose channels with 600-800um in width were used to match the size of third instar larvae. The channel is covered with a cover glass and flipped such that the larva is imaged through the glass. Representative images (overlay of brightfield and *dei^ChO^-GFP* fluorescence) show a larval ChO in a contracted **(a’)** and relaxed **(a’’)** state in the channel. **(b’-c’)**: The ChO are shown at maximal contraction for a *wt* larva (b’) and larva with altered mechanics/altered ECM composition (*prc*-mutant) (c’). The cap and ligament cells are visualized by GFP expression. Scale bar is 50um. **(b-c):** The change in the contour length of the cap cell (L_cap_) during muscle contraction as a function of R_cap_ (the distance between the endpoints of the cap cell) for a *wt* larva (b) and for a *prc*-mutant (c) with slopes of η=1.002 and η=0.27 respectively. We denote the slope by η, the efficiency, as it represents the efficiency of conversion the relative displacement of the cap cell endpoints into compressive strain within the cell. Notice that since the *wt* cap keeps its straight form during contraction, all the displacement of the cap end points is converted into a change in the cap contour length, and the slope of this curve is 1. Since the *prc* deficient cap cell experiences high-mode buckling, the slope of the curve is lower than 1 and equals ∼0.3. **(d–d’’)** Illustration of the experimental setup for assessing the femoral chordotonal organ (FCO) functionality in the adult fly. **(d)** A schematic illustration of the experimental setup used for analyzing the response of the FCO cells to joint flexion in the adult fly leg. The fly leg is dissected and mounted in a physiological medium and the FCO is visualized while femur-tibia joint flexion is induced using a mechanical probe. Representative images (overlay of brightfield and *dei^ChO^-GFP* fluorescence) of the FCO during joint extension (**d’**) and flexion (**d”**). **(e)** Representative time-course showing the relationship between the femur-tibia joint angle (green) and the FCO contour length (*L_cap_*, yellow) in *wt* leg. **(f–g)** The efficiency, η, for adult FCO *wt* with *η* = 1.03 **(f)** and *prc* mutant with *η* = 0.55 **(g).** Insets in f and g show *dei^ChO^-GFP* expression for *wt* and *prc^MB^*^03017^ FCO respectively following joint flexion. Scale bar: 100 *μ*m.

Our previous studies demonstrated that correct mechanical properties of the ChO are essential for proper function ^5^. We have shown that the loss of Pericardin (Prc), a major constituent of the ChO’s extracellular matrix^6,8^, alters the mechanical properties of the sensory organ. This change in mechanical properties led to a change in the strain (deformation) generated in the cap cell. In wildtype (*wt*) larvae, muscle-generated forces led to shortening and compressive strain generation within the cap cell with negligible bending. The compressive strain within the cap cell is most probably converted into tensile strain in the dendrite, where the mechanosensitive ion channels are located, by the unique geometrical structure of the dendritic cap as we have shown using finite-elements analysis ^5^.In contrast, in the *prc* mutants, muscle-generated forces induced significant bending in the cap cells, which underwent short-wavelength (high-mode) buckling with low compressive strain within the organ. We furthermore demonstrated that the transition from shortening (a compressed state) to bending of the cap cell interferes with its ability to propagate muscle-generated deformations correctly to the neuron and hence interferes with proper sensing ^5^ .

Here we show, in an *in-vivo* setup, that there is a critical ‘mechanical’ time window in which the ChOs must be exposed to appropriate mechanical forces in order to develop the correct mechanical properties required for organ function. Using *prc-Gal4* to drive *prc*-specific RNAi expression, we downregulated Prc at different time points during embryonic and larval life and measured the efficiency of conversion of muscle-generated mechanical deformations into compressive strain in the cap cell. When we downregulated *prc* at a late embryonic stage, or in the first or second instar larval stages, the conversion efficiency of muscle-generated deformations was reduced, however, the effect was most dramatic when *prc* was downregulated during late embryogenesis. This time window correlates with the initiation of muscle contractions in the embryo, suggesting that mechanical cues from muscles may play a critical role in the development of functional ChOs (with correct mechanical properties). Indeed, we show that reducing muscle-generated forces by a factor of 3 during the same time window, lead to the same effect as downregulating *prc* and results in abnormal sensory organ mechanics and function.

## Results

### Sensory organ function is compromised in larvae and adult flies with modified mechanical properties

ChO-specific fluorescent markers inserted into the fly genome allow us to quantify changes in the sensory organ shape during muscle contractions in live crawling larvae in a linear agarose channel using spinning disc confocal microscopy (Figure 1a and ^5,7^). In *wt* larvae, muscle contractions lead to shortening of the cap cell, where most of the change in length is accommodated. Plotting the change in the contour length of the cap cell (ΔL_cap_) during muscle contraction as a function of the distance between the two end points of the cap cell (ΔR_cap_) yields a linear relationship. For *wt* larvae, the cap cell keeps its straight form during contraction; all of the displacement of the cap cell end points is converted into a change in the cap cell contour length and therefore the slope of this curve is ∼1 (Figure 1b-b’, Video S1). In contrast, the mechanical properties of a *prc* mutant cap cell are altered and therefore it experiences high-mode buckling and the slope of this curve is lower than 1 and equals ∼0.3 (Figure 1c-c’, Video S2). We denote the slope of the curve of ΔL_cap_ vs. ΔR_cap_ by η, where η represents the efficiency of conversion of the relative displacement of the cap cell endpoints into compressive strain within the cell. For *wt* and *prc* cap cells, the average values of η are η̅ =1.01±0.01 and η̅ = 0.33 ± 0.03 respectively (Figure 2b). We have previously shown that a change in the mechanical properties of the sensory organ led to abnormal calcium firing pattern and larval locomotion in the *prc* mutant ^5^.

**FIGURE 2:**
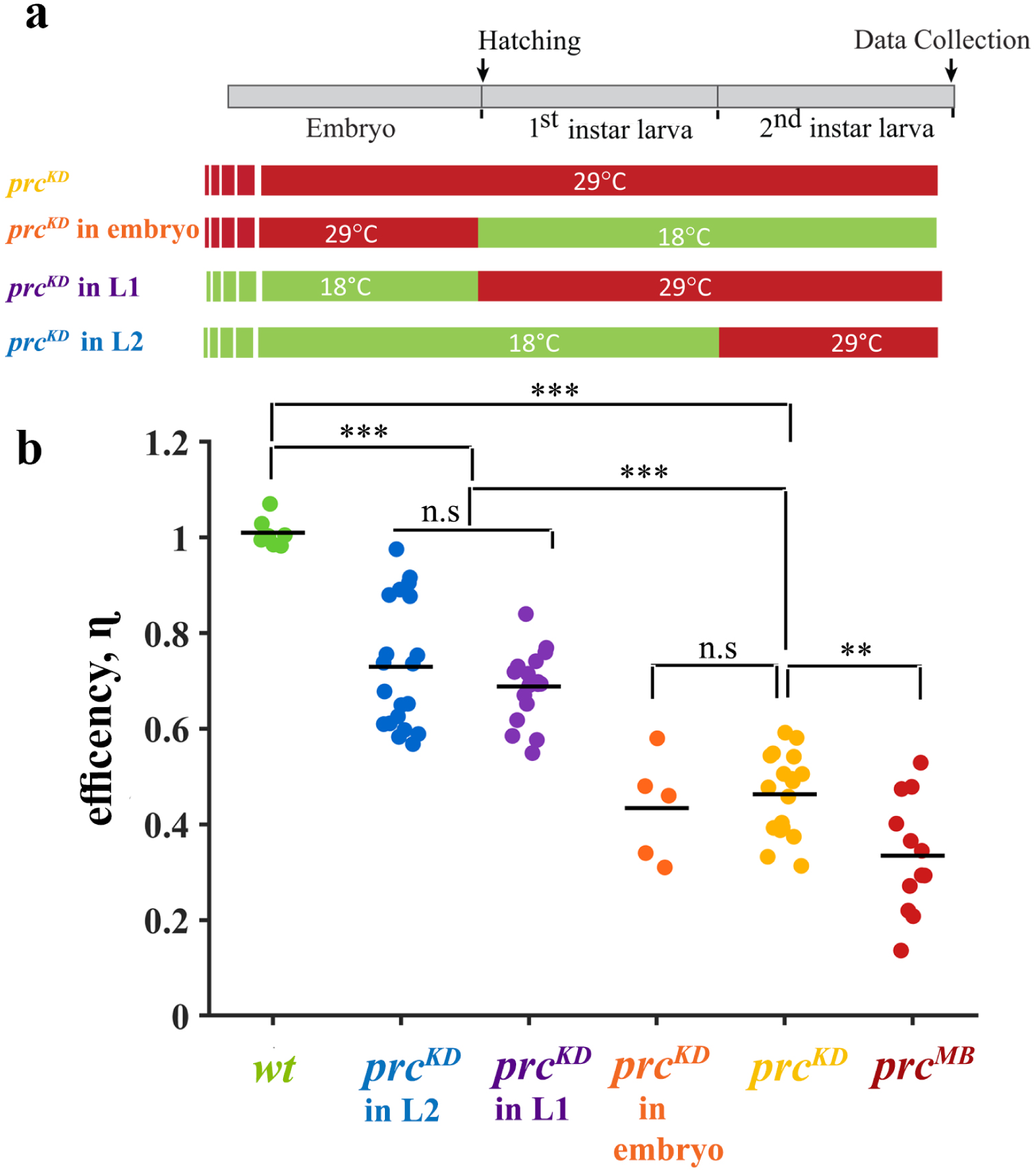
Prc-expression is required for proper development in a critical mechanical time window during embryogenesis: **(a)** Schematic illustration of the experimental protocol for time-specific reduction of *prc* expression. All experiments utilized *UAS-prc-RNAi; dei^ChO^ GFP* flies crossed to *prc-Gal4; tub-Gal80ᵗˢ*. At 18^∘^C, Gal80ᵗˢ is active and inhibits the Gal4 driver, allowing endogenous *prc* (green). At 29^∘^C, Gal80ᵗˢ is inactivated, triggering RNAi-mediated knockdown (red). Knockdown was induced at specific developmental windows and time points by shifting larvae between temperatures. **(b) :** The efficiency, *η*, for larvae in which *prc* was downregulated at different time points using different experimental protocols (shown schematically in (a)). RNAi-mediated downregulation throughout embryonic and larval development (yellow, n=18, η̅ =0.46±0.02), embryonic stage only (orange, n=5 η̅=0.43±0.05), 1^st^ instar larvae (purple, n=17 η̅=0.69±0.02) and 2^nd^ instar larvae (blue, n=20, η̅ =0.73±0.03). For comparison, we added the data for the wild-type (*wt*, green, n=7, η̅ =1.01±0.01), and *prc^MB^*^03017^ (*prc^MB^*) null mutants (red, n=12, η̅ = 0.33 ± 0.03). ***p<0.001, **p<0.01, ns=not significant. P-values were determined according to the unpaired one-tailed Mann-Whitney test.

To test whether these findings are relevant for the development of functional chordotonal organs in the adult fly, we developed a new assay that allows us to monitor the shape change of the adult fly femoral ChO (FCO) during joint flexion. Joint flexion was induced by pushing the tibia using a mechanical probe (Figure 1d-e). When changing the joint angle mechanically, we made sure that the speed of joint flexion (femur-tibia angle change) mimics the joint flexion speed observed in a walking fly^9^. Similarly to the results obtained in the larvae, while the cap cell is compressed during joint flexion in the *wt* leg (Figure 1f and Video S3), high-mode buckling is observed for the *prc* mutant leg (Figure 1g and Video S4).

### Prc expression is required for proper development in a critical time window during embryogenesis

The fly model enables temporal knockdown of *prc* expression using RNA interference (RNAi). Temporal control is achieved by incorporating *Gal80^ts^*, a temperature-sensitive inhibitor of *Gal4,* into the *Gal4/UAS* system. Optimal Gal4 activity in this system is achieved at 29°C, leading to maximum *prc* knockdown. At 18°C the Gal80^ts^ inhibits Gal4 allowing normal *prc* expression. By shifting between the restrictive (29°C) and the permissive (18°C) temperatures, the timing of *prc* downregulation can be controlled. On top of the *Gal4*/*Gal80*/RNAi components, we incorporate into the fly genome proprioceptor-specific fluorescent markers that allow us to measure the efficiency of conversion of muscle-generated mechanical deformations into compressive strain in the cap cell. To determine whether *prc* is required during a specific developmental time window, we downregulated *prc* expression in different time regimes during embryogenesis and the three instar larval stages.

To validate the effectiveness of the *prc*-specific RNAi transgene, we first downregulated *prc* throughout embryonic and larval development. This resulted in a behavior comparable to that of the null *prc* mutant (η̅ =0.46±0.02 vs. η̅ =0.33±0.03 respectively; Figure 2b) and significantly different than the *wt* (η̅ =0.46±0.02 vs. η̅ =1.01±0.01 respectively; Figure 2b). Surprisingly, when *prc* was downregulated *only* during embryogenesis, by growing the embryos in 29°C to inhibit *prc* expression and shifting larvae to 18°C immediately after hatching to restore expression in larval stages, we obtained an identical effect ( η̅=0.43±0.05; see Figure 2b).

RNAi-mediated downregulation of *prc* in the first or second instar larval stages had a smaller effect on the conversion efficiency of muscle-generated deformations into compressive strain ( η̅=0.69±0.02 and η̅ =0.73±0.03 respectively). Therefore, the full and most dramatic effect was obtained when *prc* was downregulated during late embryogenesis. This effect could not be rescued by restoring *prc* expression later during larval life. These results suggest that for the development of fully functional ChOs, *prc* expression is required within a defined critical window during embryonic development.

### Muscle-generated forces are essential for proprioceptor development

The time window in which *prc* expression is critical for proper ChO development coincides with the onset of muscle contractions in the embryo, pointing to a critical role of muscle-generated mechanical cues in the development of functional ChOs.

To test this hypothesis, we reduced muscle contractility and tested its effect on ChO development. Muscle contractility was reduced by downregulating myosin heavy chain (MHC) using RNAi. Complete elimination of MHC activity during embryogenesis is lethal. To avoid embryonic lethality and enable temporal control, we use the *Gal4*/*Gal80*/RNAi system described above. We have looked for conditions in which the level of MHC knockdown is not lethal but still significantly reduces muscle contractions. Optimal conditions were found at 27°C. Spatial specificity was achieved by inducing expression of the RNAi transgenes under the regulation of a mesoderm-specific *Gal4* driver (*mef2-Gal4*), which is not expressed in the ChO lineage. To test the hypothesis that mechanical forces are required for ChO development specifically in late embryogenesis, we downregulate MHC either throughout embryonic and larval development, or only during embryogenesis.

To quantify force reduction in the critical time window (during embryogenesis), *wt* and *mhc^KD^* embryos, both grown at 27°C, were dechorionated at stage 16 and mounted on a polyacrylamide gel (Figure 3a). Fluorescent beads embedded within the polyacrylamide gel were tracked over time and the strain field generated by each embryo was quantified (Video S5).

**FIGURE 3:**
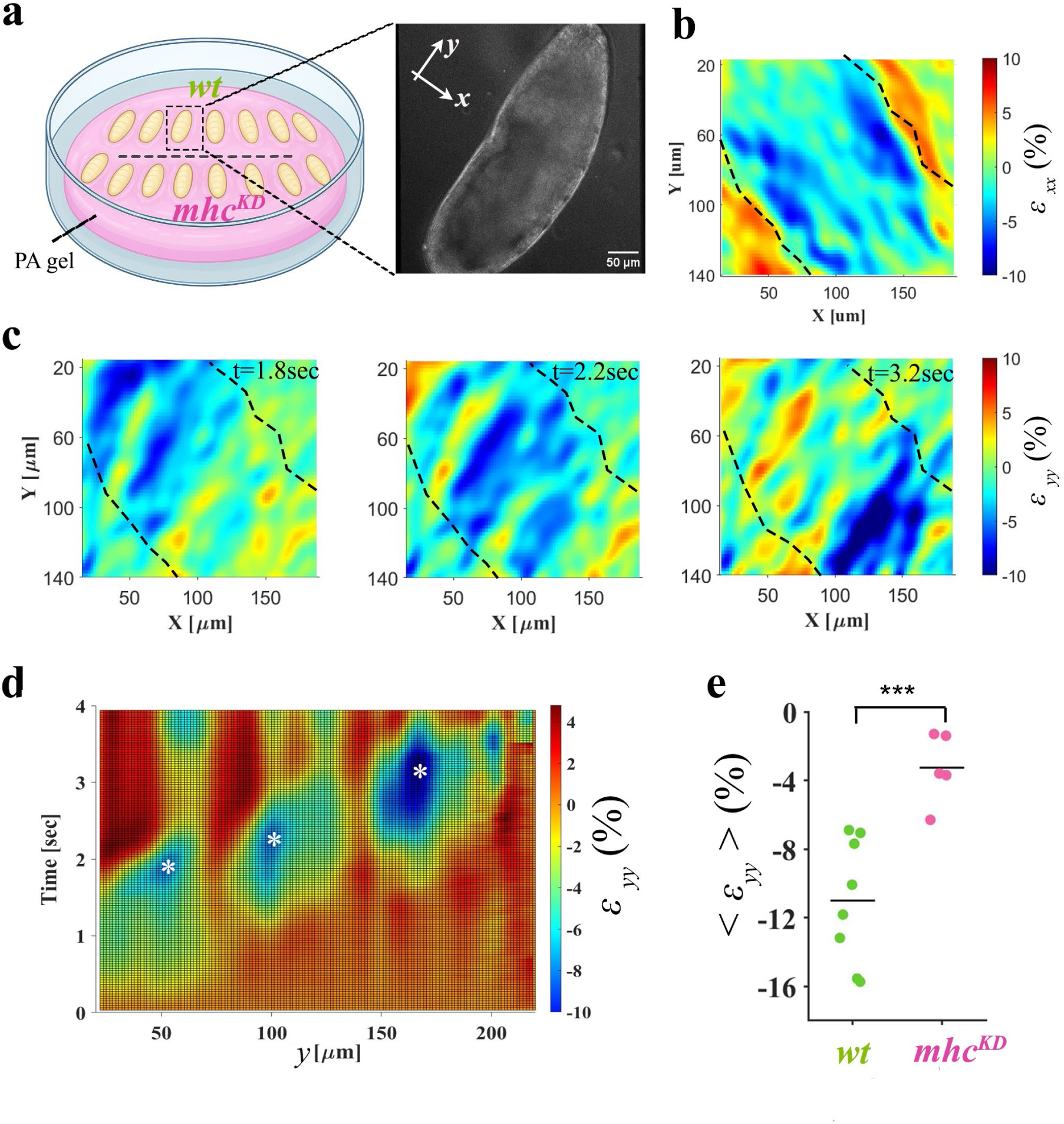
Characterization of the deformations generated by embryos with RNAi-mediated downregulation of MHC: **(a):** Schematic of the micro-mechanical sensing assay. *Drosophila* embryos are dechorionated and mounted on a compliant polyacrylamide (PA) gel substrate with 0.2*μ*m fluorescent beads (VideoS5). The inset shows a representative phase contrast image of an embryo. The lowercase x-y coordinate system denotes the analysis coordinate system: y is aligned with the direction of peristaltic wave propagation, and x is perpendicular to it. Uppercase X-Y denote the image-coordinate axes. **(b-c):** Representative strain field maps for the strain **(b)** perpendicular (ε_xx_) and **(c)** parallel (ε_yy_) to the direction of peristaltic wave propagation. Negative and positive values correspond to compression and extension respectively. Notice that the region of contraction travels as the peristaltic wave advances (VideoS5). Dashed lines indicate the embryo boundary. **(d):** Kymograph corresponding to the peristaltic wave shown in **(c).** Three contraction events for the three segments in the field of view are marked by asterisks. The average value of the three contraction events corresponds to a single point in **(e)**. **(e):** The average compressive strain, ε_yy_ for the different segments during the peristatic wave in *wt* (green, 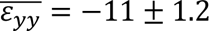) and mhc^KD^ (purple, 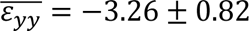). Downregulation was achieved by growing the larvae expressing the *Gal4*/*Gal80*/RNAi system at 27°C. *mhc*^KD^ larvae showed a 3.4-fold reduction in contractile force compared to wt larvae. ***p<0.001. P-values were determined according to the unpaired one-tailed Mann-Whitney test.

Drosophila larvae crawl via peristaltic waves of muscle contractions that propagate from tail to head. Muscle-generated forces start to appear around stage 16^10^ of embryonic development. Early movements are weak, local contractions, involving single segments or part of segments. These contractions increase over time and towards the end of stage 17 of embryonic development, become coordinated movements extending along the entire length of the body. Shortly before hatching full periodic peristaltic waves emerge. Only mature peristaltic waves detected shortly before hatching were considered in the analysis.

We defined the y-axis as the direction of peristaltic wave propagation and the x-axis as the perpendicular direction (Figure 3a). Figure 3b-c shows representative deformation fields parallel and perpendicular to the direction of peristaltic wave propagation. These data clearly show that the region of contraction travels as the peristaltic wave advances. Figure 3d shows a kymograph that corresponds to the peristaltic wave shown in Figure 3c. Three contraction events for the three segments in the field of view are marked by asterisks. The average compressive strain, ε_yy_ generated during the peristaltic wave by MHC-RNAi embryos (*mhc*^KD^) was 3.4-fold lower than that generated by the *wt* embryos. Since polyacrylamide behaves as a linear elastic polymer, the ratio between ε_yy_ of the *wt* embryos and that generated by the *mhc*^KD^ embryos is equal to the ratio between the contractile forces generated by embryos of these two genotypes. Thus, the contractile forces generated by the *mhc*^KD^ embryos are 3.4-fold weaker than those generated by the *wt* embryos.

We next asked whether this reduction in muscle-generated force is accompanied by altered cap cell mechanical properties and impaired proprioceptor functionality.

A change in the mechanical properties of cap cells in larvae with weakened muscles is supported by the observation that *mhc^KD^* and *wt* cap cells responded differently to the application of oscillatory local mechanical forces using a tungsten probe in a live preparation of dissected larvae (larval fillet, Videos S6-S7). The *wt* cap cell behaved mechanically like a ‘chord’ i.e., strain was generated along its entire length, and the relaxation was instantaneous (VideoS6). In contrast, the response of the *mhc^KD^* larvae was local and implies reduced bending rigidity (VideoS7), similar to the response observed for the *prc* mutant cap cell^5^

The functionality of the ChOs was assessed, as detailed above, by following the response of the ChOs to forces generated by muscle contraction in larvae crawling in a linear agarose channel (Figure 1a). As in *prc* mutant, muscle-generated forces in a *mhc*^KD^ larva induced cap cell buckling (VideoS8). As shown in Figure 4, The efficiency, η of converting muscle-generated deformation into compressive strain within the ChO cap cell for *mhc*^KD^ is reduced significantly relative to the *wt* (η̅=0.77±0.04 and η̅ =1.01±0.01 respectively). An identical effect was observed when *mhc* was downregulated *only* during embryogenesis, by growing the embryos in 27°C and shifting larvae to 18°C immediately after hatching to restore expression in larval stages (η̅=0.77±0.04; see Figure 4b). This result, consistent with our earlier finding for *prc,* further shows that restoring *mhc* expression after hatching does not rescue the phenotype caused by the lack of mechanical forces during the embryonic period.

**FIGURE 4:**
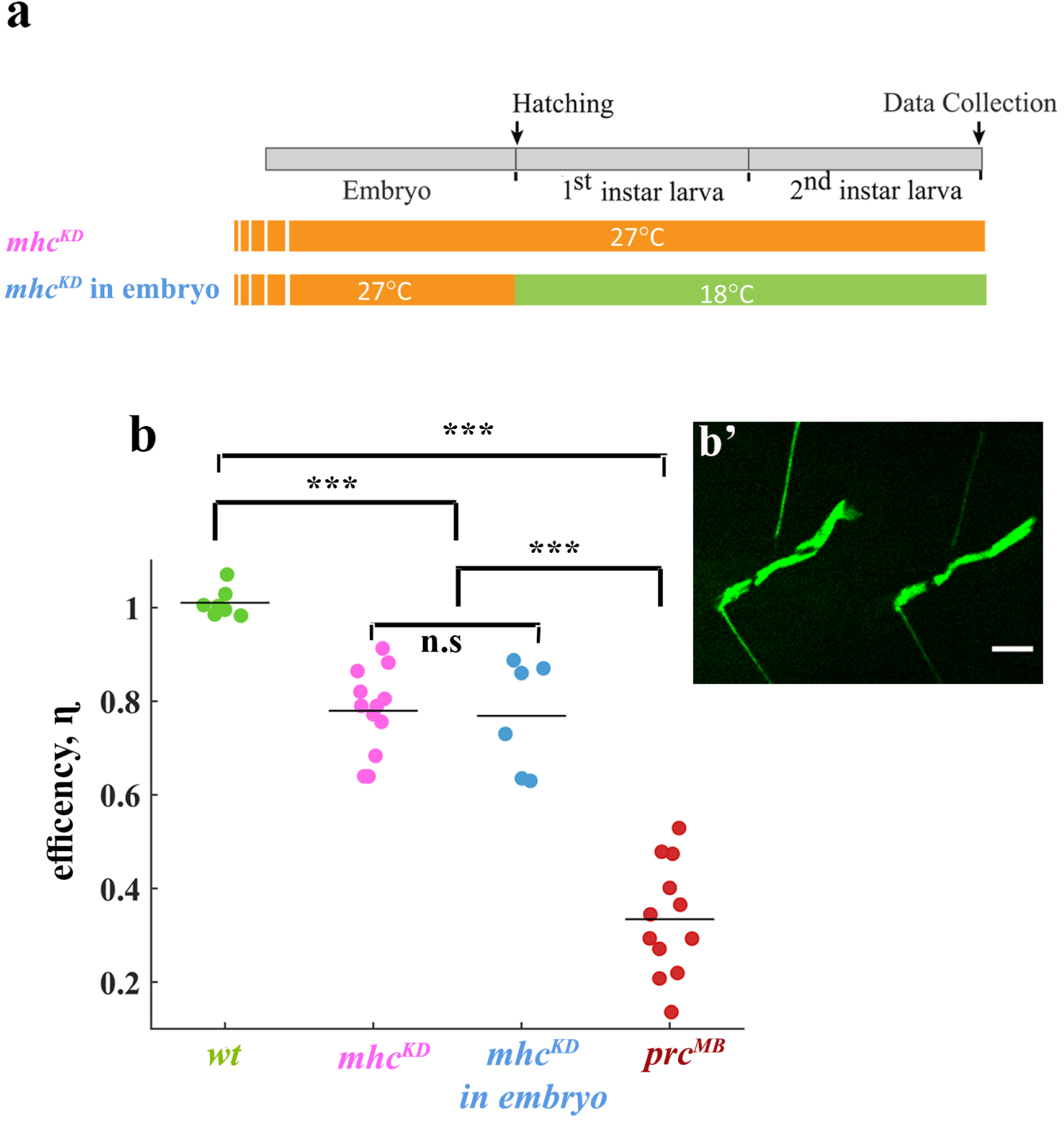
Muscle-generated forces are required for proprioceptor development: **(a) :** Schematic illustration of the experimental protocols. All experiments utilized *UAS-mhc-RNAi; dei^ChO^ GFP* flies crossed to *mef2-Gal4; tub-Gal80ᵗˢ*. Two experimental conditions are shown. *mhc^KD^* (purple): larvae were grown at 27°C throughout development (purple), resulting in partial Gal80 inactivation triggering RNAi-mediated *mhc* knowckdown that is not lethal but still significantly reduces muscle contraction (see Figure 3e). *mhc^KD^ in embryo (blue):* larvae were grown at 27°C only during the embryonic period and then shifted to 18°C after hatching. At 18^∘^C, Gal80ᵗˢ is active and inhibits the Gal4 driver, allowing endogenous *mhc* expression. **(b) :** The efficiency, η of converting muscle-generated deformation into compressive strain within the ChO cap cell using the experimental protocols shown schematically in (a). Data are shown for *mhc^KD^* (purple, n=12, η̅ =0.78±0.02) and *mhc^KD^* in embryonic stage only (blue, n=6, η̅ =0.77±0.04). For comparison, we show again the data from Figure 2b for wild type (*wt*, green, n=7, η̅ =1.01±0.01) and *prc^MB^*null mutants (red, n=12, η̅ =0.33±0.03). **(b’):** Spinning disc confocal images of ChO shape during muscle contraction in *mhc^KD^*. The cap and ligament cells are visualized by GFP expression. Muscle contractions induce cap cell buckling in *mhc^KD^* similar to the phenotype obtained for *prc^KD^*. Scale bar: 50 *μ*m. ***p<0.001, ns=not significant. P-values were determined according to the unpaired one-tailed Mann-Whitney test.

## Discussion

Using the ChO of the fly larva as a model system, we show in an *in-vivo* setup that mechanical forces are required for proprioceptor development with correct mechanical properties and sensory function. We have previously shown^5^ that eliminating Prc, a major ECM component of the ChO, alters the mechanical properties of the sensory organ. We further showed that this change leads to short-wavelength buckling of the cap cell upon muscle contraction with low compressive strain within the cell which leads to impaired sensory function. By reducing Prc expression at different developmental stages, we show here that there is a critical time window where Prc expression is essential for the development of a sensory organ with proper mechanical properties that allow sufficient compressive strain to build up in the cap cell and thereby enable normal sensory function. We refer to this period as a ‘mechanical time window’ since it coincides with the onset of mechanical forces in the embryo. Consistent with this idea, reducing the forces generated by the muscles in the embryo results in a very similar phenotype. Since full knockout of *mhc* activity is lethal, we were not able to characterize the effect of complete elimination of muscle-generated forces on ChO development. Nevertheless, we were able to show that even by reducing muscle-generated forces during embryogenesis by 3.4-fold, we obtained a significant phenotype that cannot be rescued by expressing normal levels of MHC later during larval life.

The fact that the *prc*-specific RNAi and *mhc*-specific RNAi induce very similar effects on ChO behavior, and that in both cases the strength of the phenotype depends similarly on the developmental timing of knockdown, points to a mechanism whereby Prc is crucial for correct mechanotransduction. This can occur, for example, if Prc, due to its self-assembly properties acts as a scaffold for the formation of adhesions with the correct composition and size, thereby influencing mechanotransduction.

The cytoskeleton of the ChO cap cell consists mainly of microtubules ^11,12^, whereas actin is present mainly at the ECM adhesion sites. It is possible that posttranslational modifications of microtubules as well as changes in microtubule crosslinker expression or a shift in the composition of tubulin isoforms^16,19^ play an important role in modifying the cap’s mechanical properties. Double mutants for tubulin/tubulin-associated proteins were shown to induce compression to buckling/supercoils transition in touch receptor neurons in *C. elegans*^20,21^.

In Drosophila, the major posttranslational modifications are acetylation, glycylation and glutamylation. Microtubule acetylation was found to protect microtubules from catastrophe and to lead to increased elasticity^15^ . It was recently shown that microtubule deacetylation can be mediated by mechanosensitive ion channels^18^ and that mechanical signaling promotes talin-mediated recruitment of αTAT1 (β-Tubulin acetylase) to focal adhesions which leads to local microtubule acetylation^13^. In any event, our data suggest that all these rearrangements must occur within an early and well-defined time window during embryogenesis.

## Supplementary information

is available.

## Acknowledgements

We thank the Bloomington Drosophila Stock Center, and the Vienna Drosophila Resource Center for fly strains. We are thankful to E. Heckscher for the kind gift of an agarose channel casting mold. This research was supported by the Israel Science Foundation (grant 1664/21) and funds from the Rappaport Family Institute for Research in the Biomedical Sciences.

## Author contributions

S.T and A.S. designed and supervised the research and wrote the manuscript. A.H. developed and performed the crawling larvae, adult fly and embryo assays, N.H constructed the required fly strains and performed fly work.

## STAR METHODS

### EXPERIMENTAL MODEL AND SUBJECT DETAILS

#### Fly strains

The *dei*^ChO^-GFP reporter strain^14^ was used as the control. The *Mi{ET1}prc^MB^*^03017^ allele was used as a *prc* null allele. For conditional knockdown of *prc*, *UAS-prc*-RNAi; *dei*^ChO^-GFP flies were crossed to *prc*-Gal4; *tub*-Gal80^ts^. Muscle-specific knockdown experiments were performed using *mef2*-Gal4; *tub*-Gal80^ts^ crossed to *UAS-mhc*-RNAi; *dei*^ChO^-GFP. All crosses were maintained at permissive temperature and shifted to the restrictive temperature at the indicated developmental stages to induce RNAi expression.

### METHOD DETAILS

#### Imaging the ChO of larvae crawling in linear agarose channels

Larvae containing the *dei^ChO^-GFP*reporter, were used for monitoring cell shape changes in the ChO cells during larval locomotion. Larvae were derived from 8-hour egg lays, collected at the first 8 hours of the third instar stage and placed in a 1cm long linear channel made of 3% agarose. The channels (600,700, or 800 μm wide) were prepared as described in ^7^, using a 3D-printed casting mold. To image the ChO, the orientation of the larva was adjusted by rolling it in the channel to ensure a lateral view. High-speed imaging was performed using a Nikon Ti-E inverted microscope equipped with a Yokogawa spinning disk confocal system. This setup enabled tracking of fast larval movement at an acquisition rate of 33 frames per second.

#### Mechanical stimulation assay

Mechanical stimulation was applied using a rigid tungsten probe with a 25μm tip diameter (Signatone) mounted on a three-axis (X–Y–Z) micromanipulator (Thorlabs) attached to a custom-built microscope stage adaptor. To enable simultaneous visualization of the probe and ChO cap cells. The probe was positioned adjacent to the center of the cap cell and oscillated perpendicular to the longitudinal axis of the cap cell with an amplitude of 25 μm using an arbitrary function generator (Stanford Research Systems DS345). A similar setup has been described previously^17^.

#### Preparation of polyacrylamide coated plates

Polyacrylamide (PA) gel coated plates were prepared following the protocol described previously^25^. Briefly, 30 mm glass-bottom plates (Greiner) were activated using 2% 3-aminopropyltrimethoxysilane, and 1% glutaraldehyde solution. Filtered polyacrylamide/bis-acrylamide 7.5/0.03% solution (Bio-Rad) was mixed with 0.02% suspension of carboxylated dark red fluospheres (Life Technologies), 0.05% ammonium persulphate and 0.4% temed (Bio-Rad). 15 μl of mixed PA solution was applied on each activated plate and covered with an 18 mm round glass cover slip to form a thin film. PA films were left to polymerize for 30 min at room temperature then covered with PBS and saved at 4C until used.

#### Live Imaging of embryos and Traction Force Microscopy

Drosophila embryos were collected for 2 hours and maintained at 27°C until early stage 17 (approximately 14 hours AEL), and dechorionated in 50% bleach for 1 minute. Following water washes, embryos were gently mounted on polyacrylamide (PAA) gels functionalized with Sulfo-SANPAH to facilitate stable adhesion. wt and mutant embryos were put on the same gel substrate. To prevent desiccation, embryos were covered with Halocarbon oil (Sigma-Aldrich). Live imaging was performed using a Nikon Ti-E inverted microscope equipped with a Yokogawa spinning disk confocal system. Fluorescent beads embedded within the PAA gel were simultaneously imaged using a 561 nm laser line. These bead displacements served as the basis for subsequent Traction Force Microscopy (TFM) and DIC-based mechanical analysis to quantify the forces exerted by the muscle contraction before embryo hatching.

### QUANTIFICATION AND STATISTICAL ANALYSIS

Statistical significance is indicated by n.s. for no significance, * for p<0.05, ** for p<0.01 and *** for p<0.001. P-values were determined according to the unpaired one-tailed Mann-Whitney test. The number of biological repeats is stated in the figure legends.

Data analysis: Basic image analysis is done using ImageJ and custom-written MATLAB codes.

