## Supplementary material for "A Mechanical time window for correct proprioceptor development": Video_Legends

**Video S1: ChO imaging during muscle contraction in a *wt* crawling larva**

Time lapse imaging of GFP-expressing ChOs during muscle contraction in a *wt* larva crawling in an agarose channel. The *wt* cap cell becomes shorter during muscle contraction, while maintaining its straight form. Scale bar is 100µm. The movie is played at real time speed.

**Video S2: ChO imaging during muscle contraction in a *prc* crawling larva**

Time lapse imaging of GFP-expressing ChOs during muscle contraction in a *prc* larva crawling in an agarose channel. The *prc* cap cell is buckled during muscle contraction. Scale bar is 100µm. The movie is played at real time speed.

**Video S3: Adult femoral ChO imaging during joint flexion in a *wt* leg**

Time lapse imaging of GFP-expressing FCOs during joint flexion in a *wt* leg. Joint flexion was induced by pushing the tibia using a mechanical probe. *left*: magnification of 40x (scale bar is 50µm) *right:* magnification of 10x (scale bar is 100µm). It is clearly shown that the cap cell is compressed maintaining its straight form during joint flexion. The movie is played in real time speed.

**Video S4: Adult femoral ChO imaging during joint flexion in a *prc* mutant leg**

Time lapse imaging of GFP-expressing FCOs during joint flexion in a *prc* leg. Joint flexion was induced by pushing the tibia using a mechanical probe. *left*: magnification of 40x (scale bar is 50µm). *right*: magnification of 10x (scale bar is 100µm). It is clearly shown that the *prc* cap cell is buckled during joint flexion. The movie is played in real time speed.

**Video S5: Strain field generated by muscle-generated forces in Drosophila embryo**

*(left)*: Time lapse imaging of the deformation field generated by muscle-generated forces of a Drosophila embryo. Mechanical deformations are detected by following movements of 0.2µm fluorescent beads embedded in the gel. Scale bar is 50µm.

The corresponding strain maps are shown on the center and the right along the direction of the peristaltic wave ($\varepsilon_{yy}$) and in the perpendicular direction ($\varepsilon_{xx}$) respectively. The video is played in real time.

**Video S6: *wt* cap cell response to application of oscillatory mechanical force**

Time lapse movie of *wt* cap cell response to the application of oscillatory mechanical deformations by a tungsten probe. The cap and ligament cells are visualized by GFP expression. Scale bar: 50µm. The movie is played in real time speed.

**Video S7: *mhc$^{KD}$* cap cell response to the application of oscillatory mechanical force**

Time lapse movie of *mhc$^{KD}$* cap cell response to the application of oscillatory mechanical deformations by a tungsten probe. The cap and ligament cells are visualized by GFP expression. Scale bar is 50µm. The movie is played in real time speed.

**Video S8: ChO imaging during muscle contraction in a *mhc$^{KD}$* crawling larva**

Time lapse imaging of GFP-expressing ChOs during muscle contraction in a *mhc$^{KD}$* larva crawling in an agarose channel. The *mhc$^{KD}$* cap cell is buckled during muscle contraction. Scale bar is 100µm. The movie is played at real time speed.
